# Distinct Structural Flexibility within SARS-CoV-2 Spike Protein Reveals Potential Therapeutic Targets

**DOI:** 10.1101/2020.04.17.047548

**Authors:** Serena H. Chen, M. Todd Young, John Gounley, Christopher Stanley, Debsindhu Bhowmik

## Abstract

The emergence and rapid worldwide spread of the novel coronavirus disease, COVID-19, has prompted concerted efforts to find successful treatments. The causative virus, severe acute respiratory syndrome coronavirus 2 (SARS-CoV-2), uses its spike (S) protein to gain entry into host cells. Therefore, the S protein presents a viable target to develop a directed therapy. Here, we deployed an integrated artificial intelligence with molecular dynamics simulation approach to provide new details of the S protein structure. Based on a comprehensive structural analysis of S proteins from SARS-CoV-2 and previous human coronaviruses, we found that the protomer state of S proteins is structurally flexible. Without the presence of a stabilizing beta sheet from another protomer chain, two regions in the S2 domain and the hinge connecting the S1 and S2 subunits lose their secondary structures. Interestingly, the region in the S2 domain was previously identified as an immunodominant site in the SARS-CoV-1 S protein. We anticipate that the molecular details elucidated here will assist in effective therapeutic development for COVID-19.

## I. Introduction

A new coronavirus, severe acute respiratory syndrome coronavirus 2 (SARS-CoV-2), causes respiratory illness and has now reached a pandemic scale. Denoted as coronavirus disease 2019 (COVID-19), its global spread is currently ongoing with symptoms that can range from mild flu-like to severe pneumonia, leading to death in certain cases. Early investigations in China showed that SARS-CoV-2 has a high genomic sequence similarity to the previous SARS-CoV-1, along with a bat coronavirus [1], [2]. Similar to SARS-CoV-1, SARS-CoV-2 is a positive-sense, single-stranded RNA virus of the betacoronavirus genus.

Given the health crisis caused by the mounting number of COVID-19 cases worldwide, there is an urgent need to develop effective therapeutics and eventual vaccines. A critical step in developing targeted treatments is obtaining a detailed understanding of the molecular pathways of SARS-CoV-2 and constituent structures. The structural biology community has made rapid progress towards this goal by experimentally determining several SARS-CoV-2 proteins, including the spike (S) [3]–[6], nucleocapsid (N) [7], and main protease (M^pro^) [8]. The S protein is particularly important since it resides on the viral envelope and is responsible for host cell entry by engaging angiotensin-converting enzyme 2 (ACE2) receptors [2], [9], [10]. Recent experimental structures of the SARS-CoV-2 S protein receptor binding domain (RBD) in complex with ACE2 provide detailed interface information [4], [6]; targeting this interface represents an active area of research for therapeutic development [11]. However, there may exist potential targets on the S protein besides the RBD domain.

Further motivating our work is the need to understand SARS-CoV-2 structure in the context of structures from previous coronaviruses. Prior to the emergence of SARS-CoV-2, comparisons of the trimeric S protein from different viruses showed they possess overall similar structures but with some local differences [12], [13]. With the appearance of SARS-CoV-2, it is necessary to investigate the S protein structure [3], and compare it to previous human coronaviruses.

Here, to obtain deeper insights into S protein structure for biological understanding and therapeutic targeting, we employ a combined molecular dynamics (MD) simulation and artificial intelligence (AI) methodology on a series of coronavirus S proteins. Specifically, we investigate the S proteins from the current SARS-CoV-2, SARS-CoV-1, Middle East respiratory syndrome coronavirus (MERS-CoV), and human coronavirus HKU1 (HCoV-HKU1). From structural analysis of extensive MD simulations, we find substantial flexibility between the subunits of the S proteins. We find that reduced distance matrix representations, interpreted by unsupervised deep learning (DL), reveal important regions for S protein trimerization. These regions present potential targets for therapeutic development.

## II. Results and Discussion

Human coronavirus spike (S) proteins are molecular complexes each formed by three protein chains [3], [14]–[16]. To better understand S protein structure [3], we started by studying the protomer, the structural unit of the trimeric complex, and compared the protomer structure of the current SARS-CoV-2 with protomers of previously identified human coronaviruses, including SARS-CoV-1 [15], MERS-CoV [16], and HCoV-HKU1 [14]. We modeled each protomer system from the corresponding cryo-EM S protein structure (see Methods for more details). There are three major structural domains which constitute the protomer: the amino-terminal domain (NTD), receptor binding domain (RBD), and the S2 domain.

The NTD and RBD are within the S1 subunit and are responsible for binding to host receptors, while the S2 domain is in the S2 subunit which mediates fusion of the viral and host membranes [14], [15]. Fig. 1 A-D shows the initial protomer structures of the four S proteins, each highlighted by the three domains. To determine the dynamic change of the domain organization in the protomer structures, we measured the distribution of the interdomain distances from 5,000 structures of each system taken from 25 independent 200 ns MD simulations (Fig. 1 E-G). Given the limited timescale, we made no attempt to reproduce the thermodynamics but focused on computing the structural features of the systems [17]. The overall distributions of the three interdomain distances are broad, ranging from 40 Å to 110 Å between S2 and NTD/RBD domains and 30 Å to 100 Å between NTD and RBD domains. These wide distributions indicate enhanced structural flexibility in the domain arrangement of the four protomer systems compared to the cryo-EM structures.

**Figure 1.**
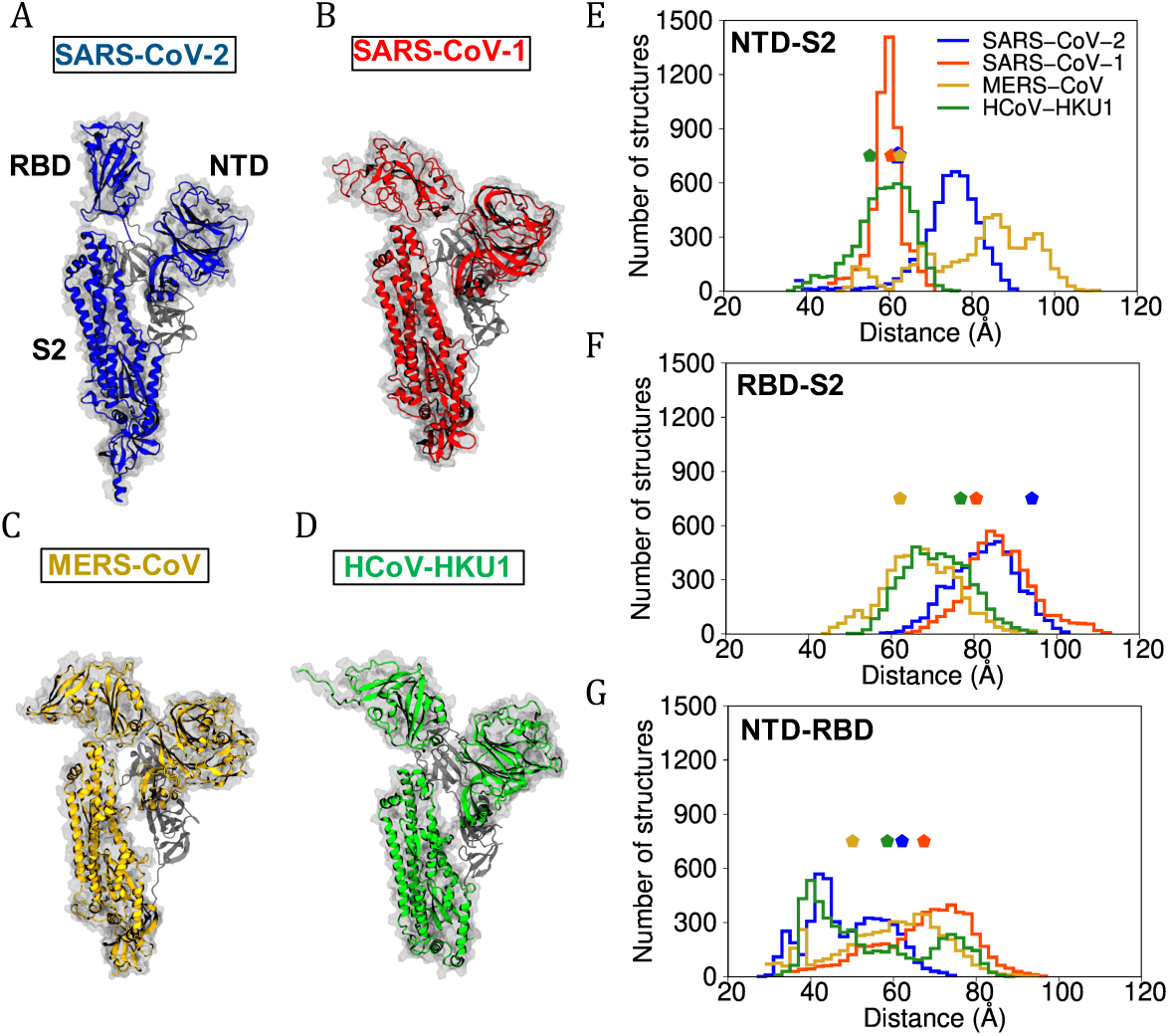
Structure and distribution of the interdomain distances of the four spike protein protomers. (A-D) The initial protomer structure of (A) SARS-CoV-2, (B) SARS-CoV-1, (C) MERS-CoV, and (D) HCoV-HKU1. The three major domains, NTD, RBD, and S2, are labeled. (E-G) The distribution of the center-of-mass distance between the backbone atoms of the residues that constitute (E) NTD and S2, (F) RBD and S2, and (G) NTD and RBD calculated from 5,000 structures of each system taken from the MD simulations. The distribution of the SARS-CoV-2 protomer is colored in blue, SARS-CoV-1 in red, MERS-CoV in yellow, and HCoV-HKU1 in green. The distances calculated from the cryo-EM structures are marked by a pentagon.

To gain further insight into the 20,000 protomer structures, we applied convolutional variational autoencoders (CVAEs) to encode the high dimensional protein structures from the MD simulations into 3-D latent spaces for visualization. Compared to commonly applied structural analysis methods, which are mostly limited to specific regions of the protein of interest, our DL approach allows us to evaluate a protein structure as a whole yet still capture detailed local structural features. We found individual clusters corresponding to each of the four protomer systems, suggesting that the structural features of the four systems embedded in the latent spaces are distinct from each other (Fig. 2 A-C). Representative structures selected from the clusters also show high structural flexibility in their domain arrangement (Fig. 2 D-F). The hinge region connecting the S1 and S2 subunits opens up, causing broad distributions of the interdomain distances observed in Fig. 1 E-G.

**Figure 2.**
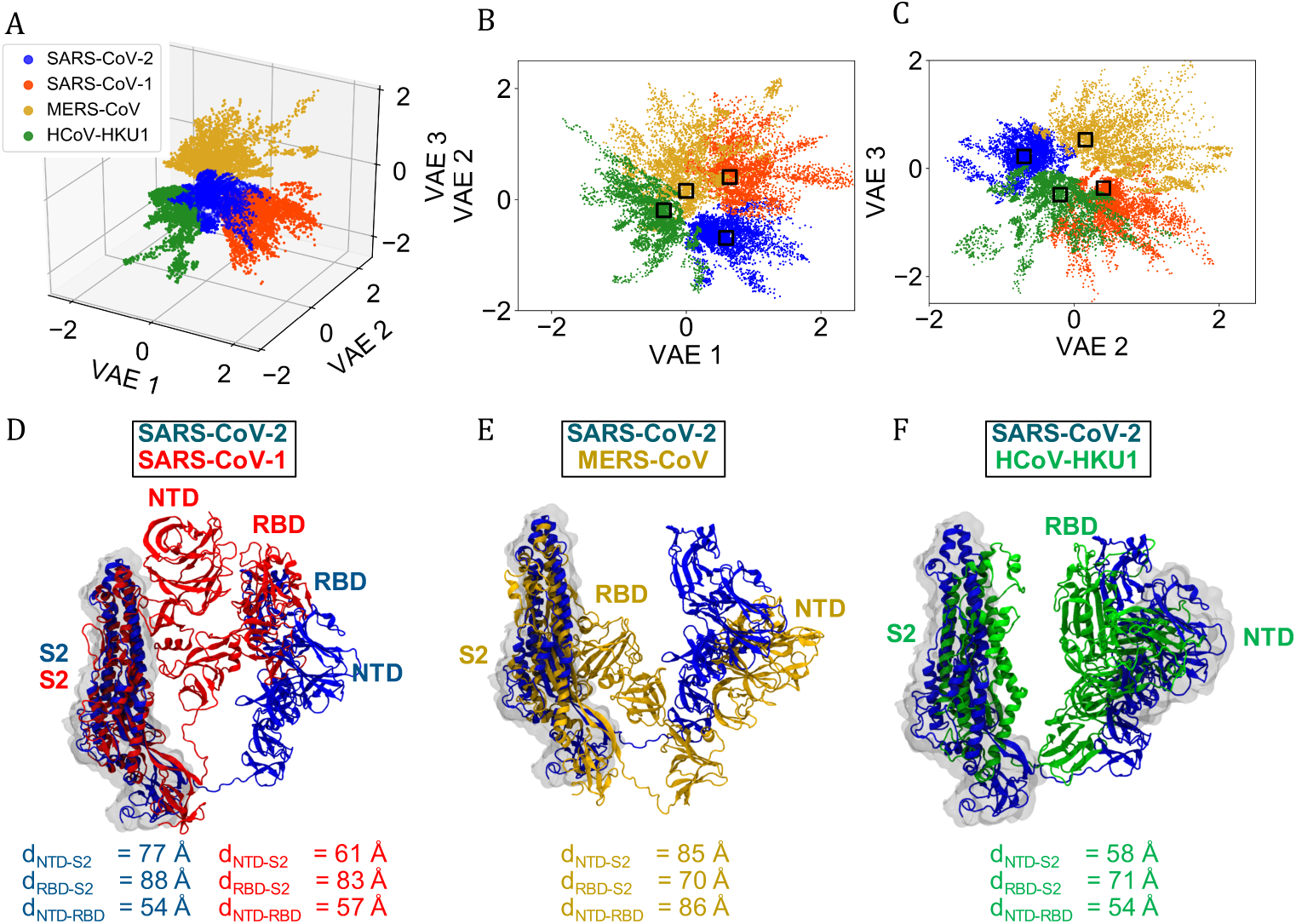
Structural profiles of the four spike protein protomers. (A) The 3-D latent space representing a total of 20,000 structures labeled by the virus system name. The structures of the SARS-CoV-2 protomer is colored in blue, SARS-CoV-1 in red, MERS-CoV in yellow, and HCoV-HKU1 in green. (B) The projection of (A) onto the VAE 1/ VAE 2-plane. (C) The projection of (A) onto the VAE 2/ VAE 3-plane. Four representative protomer structures are selected from the clusters, each outlined in a box in (B-C). (D-F) The four representative protomer structures shown in cartoon representation. The SARS-CoV-2 protomer structure is aligned with (D) SARS-CoV-1, (E) MERS-CoV, and (F) HCoV-HKU1, respectively. The aligned regions are in surface representation. The interdomain distances of the four structures are included.

The exposed surface in the protomer may provide new targets for therapeutic action. Therefore, we further explored the structural flexibility of the SARS-CoV-2 S protein. We compared the structures between the protomer and trimer to investigate whether the large structural arrangement of the domains in the protomer is affected by the trimeric state. We found that domain organization is stabilized by trimerization. Unlike the protomer, the three domains are closely arranged in the trimer (Figs. 3 A-B and S4). In addition to their interdomain distances, the differences in structural flexibility between the protomer and trimer is captured in solvent-accessible surface area (SASA) values (Fig. 3 C). The SASA values of the protomer and one chain of the trimer are 532 *±* 10 nm^2^ and 450 *±* 8 nm^2^, respectively. The difference in these values largely comes from their S2 domains, which have the values of 228 *±* 7 nm^2^ and 170 *±* 5 nm^2^, respectively. Again, we applied a CVAE to encode the 10,000 protomer and trimer structures into 3-D latent spaces. We found two individual clusters, one corresponding to the protomer and the other to the trimer, suggesting that the structural features of the two oligomeric states embedded in the latent spaces are different (Fig. 3 D-G).

**Figure 3.**
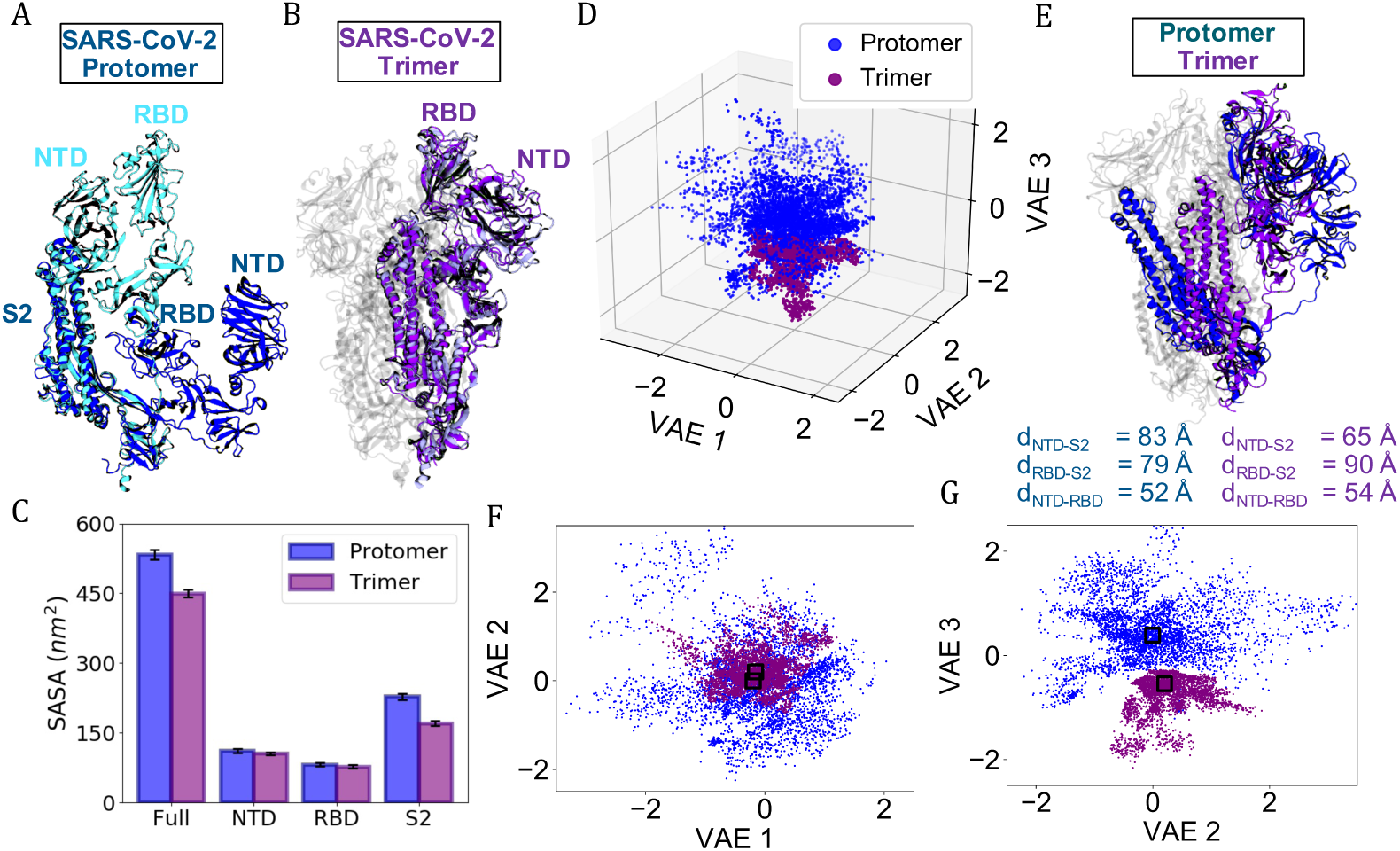
Structural profiles of the protomer and trimer of SARS-CoV-2 spike protein. (A) The first (light blue) and last (dark blue) structures from one representative MD trajectory of SARS-CoV-2 protomer. (B) The first (light purple) and last (dark purple) structures from one representative MD trajectory of SARS-CoV-2 trimer. (C) The SASA values of the full protein and three domains in the protomer and trimer. Each bar represents the average calculated from the MD structures in the corresponding 25 trajectories. The error bars represent the standard deviation. (D) The 3-D latent space representing a total of 10,000 structures labeled by the oligomeric state. (E) Representative protomer and trimer structures selected from the clusters, each outlined in a box in (F-G) and shown in cartoon representation. The interdomain distances of the structures are included. (F) The projection of (D) onto the VAE 1/ VAE 2-plane. (G) The projection of (D) onto the VAE 2/ VAE 3-plane.

To locate the different structural features between the protomer and trimer in their 3-D structures, we compared the difference between the distance matrices of the representative protomer and trimer structures selected from the clusters in the latent dimensions (Fig. 4 A). Most differences between the distance matrices result from interdomain arrangement. However, there are some disparities within the S2 domain (Fig. 4 B). One region of difference corresponds to residue numbers 784 to 810 in the PDB. In the 3-D structure, this region forms a beta sheet in the trimer while the structure is lost in the protomer (Fig. 4 C). Further investigation into this region reveals that the beta sheet in the trimer is stabilized by another beta sheet constituted by residues 700 to 710 of another chain (Fig. 4 D). Without the presence of other chains, this stability is lost; not only do residues 784 to 810 in the S2 domain lose the beta sheet structure, but residues 700 to 710 in the hinge region connecting the two subunits become an extended loop (Fig. 4 E). Oligomeric proteins are often stabilized by oligomerization and hold highly flexible regions in the protomer state [18]. The transition between the structured and unstructured form of these flexible regions is sometime reversible, but due to the size of the S protein protomer and the timescale applied in this study, we did not observe reversible behavior of the two loop regions in the protomer state. Interestingly, a fragment between residues 784 and 810 was identified previously as an immunodominant site in SARS-CoV-1 S protein [19]. Complementary antibodies acting on this site provided the dominant immune response for patients who recovered from the SARS-CoV 1 infection. Our results provide structural comprehension on the previous experimental observation and posit that targeting these residues might not only interfere with S protein trimerization and the subsequent viral activity but also aid antibody immune response. Our findings provide insights into therapeutic design targeting the S protein beyond the oft-targeted RBD domain.

**Figure 4.**
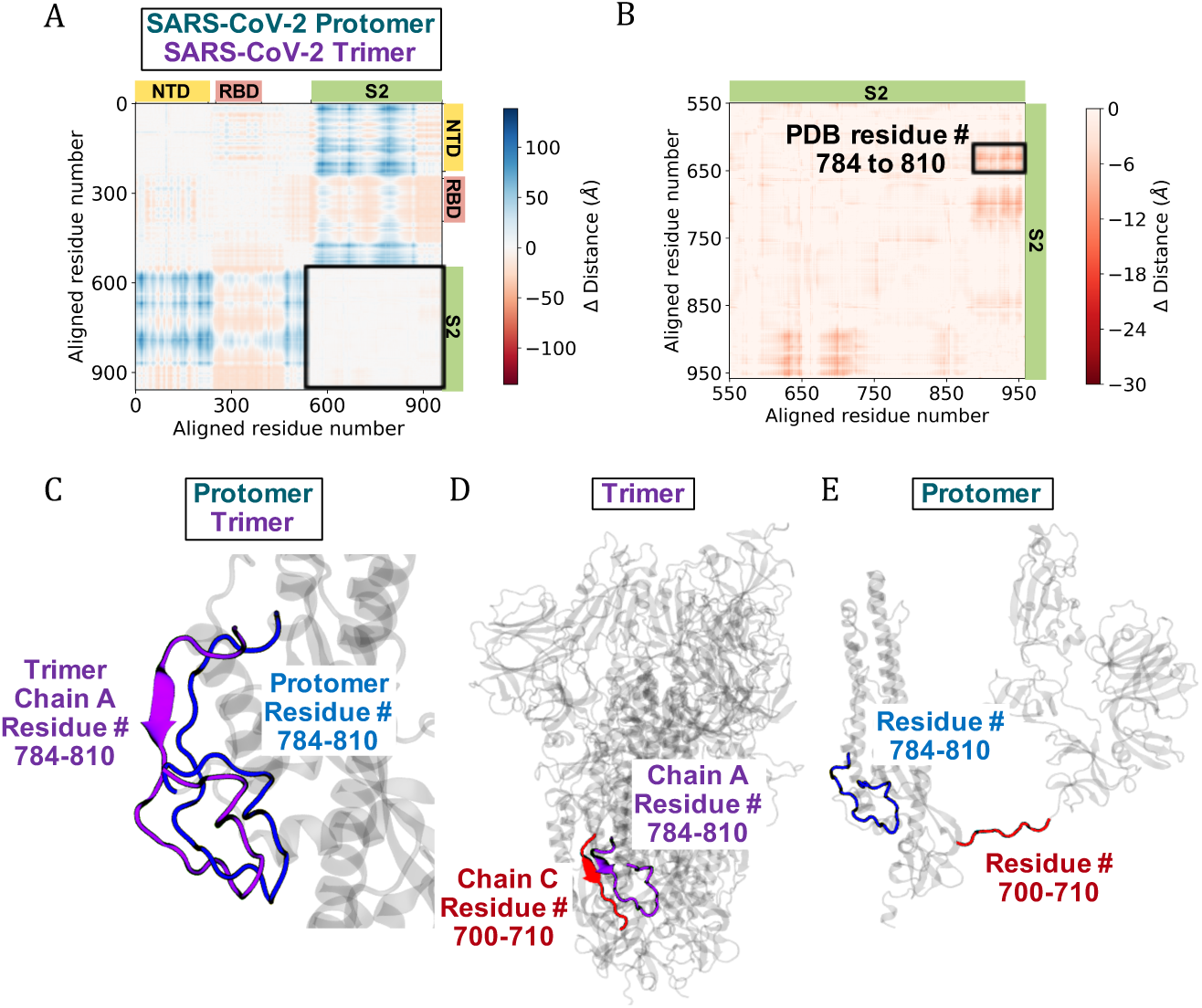
Differences in the distance matrices of the protomer and trimer of SARS-CoV-2 spike protein reveal distinct structural features in the S2 domain and the hinge region. (A) A heat map representing the differences between the distance matrices of the protomer and trimer of SARS-CoV-2 spike protein shown in Fig. 3 E. The aligned residue numbers corresponding to the three domains are labeled on the map. (B) S2 domain enlarged. The structural feature of interest is boxed and the corresponding residue numbers in the PDB are labeled. (C) Magnified Fig. 3 E to the location of the structural feature identified from the heat map. The structure from the trimer is colored in purple while the protomer is in blue. (D) Structural features identified in the trimer. The red beta sheet from a different protomer stabilizes the purple beta sheet in the trimer. (E) The same features illustrated in the protomer with the loss of the stabilizing beta sheet.

## III. Conclusion

Here, we used unsupervised deep learning routines based on CVAEs to systematically compare S protein ensembles from MD simulations across lower-dimensional, latent spaces. By first comparing the S protein protomer structure of SARS-CoV-2 to those from previous human coronaviruses, we identified distinct clusters for each virus in the 3-D latent space, where representative structures from these clusters highlight their differences in domain flexibility. Next, we compared the SARS-CoV-2 S protein protomer and trimer structures, which also displayed a clear separation of clusters in the latent space. While the main distinctions between these two states arise from the general gain in structural stability as the protomer self-assembles into the trimer state, we pinpointed structural transitions in specific flexible regions of the protomer that warrant consideration as potential therapeutic targets. These regions are promising as natural targets of immune recognition, but more importantly, they are involved in S protein oligomerization, suggesting they are susceptible to therapeutic action for protein destabilization. Overall, our study provides a more complete molecular view of the SARS-CoV-2 S protein that may assist in accelerating both vaccine and drug design efforts.

## IV. Methods

### A. Molecular systems and MD simulations

To generate initial systems for the structural study of human coronavirus S proteins, we built protomer structures from chain A of the cryo-EM S protein structures of SARS-CoV-2 (PDB 6VSB [3]), SARS-CoV-1 (PDB 6CRZ [15]), MERS-CoV (PDB 6Q05 [16]), and HCoV-HKU1 (PDB 5I08 [14]). The trimeric state of the SARS-CoV-2 S protein consists of all three chains in PDB 6VSB. Each structure was solvated in the center of a water box with a minimum distance of 15 Å from the edge of the box to the nearest protein atom, neutralized with counter ions and ionized with 150 mM NaCl. Table 1 summarizes the details of the five molecular systems studied.

**TABLE I.**
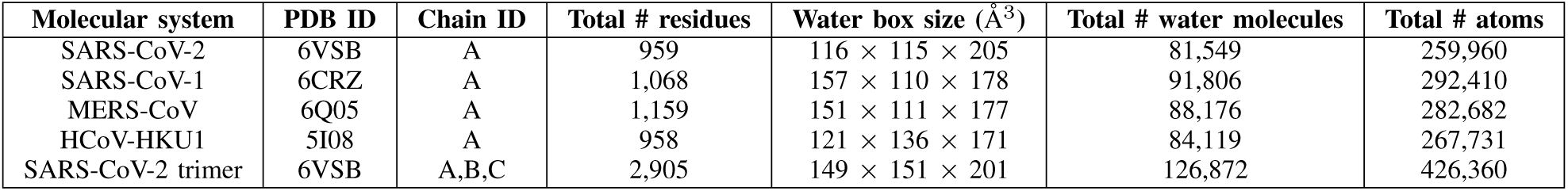
Summary of the molecular systems

| Molecular system | PDB ID | Chain ID | Total # residues | Water box size ($\text{\AA}^3$) | Total # water molecules | Total # atoms |
| --- | --- | --- | --- | --- | --- | --- |
| SARS-CoV-2 | 6VSB | A | 959 | $116 \times 115 \times 205$ | 81,549 | 259,960 |
| SARS-CoV-1 | 6CRZ | A | 1,068 | $157 \times 110 \times 178$ | 91,806 | 292,410 |
| MERS-CoV | 6Q05 | A | 1,159 | $151 \times 111 \times 177$ | 88,176 | 282,682 |
| HCoV-HKU1 | 5I08 | A | 958 | $121 \times 136 \times 171$ | 84,119 | 267,731 |
| SARS-CoV-2 trimer | 6VSB | A,B,C | 2,905 | $149 \times 151 \times 201$ | 126,872 | 426,360 |

Following a similar protocol to our previous studies [20], [21], the resulting systems were each subjected to 20,000 steps of energy minimization, followed by 1 ns equilibration with harmonic restraints placed on the heavy atoms of the protein. The force constant was 1 kcal/mol/Å [15]. After equilibration, the restraints were released, and a 200 ns trajectory was generated in a production run. For each system, 25 independent 200-ns trajectories were performed. Structures were taken every 1 ns for analysis, yielding a total of 5,000 structures for each system.

All MD simulations were performed with NAMD [22] in NPT ensemble at 1 atm and 310 K with a time step of 2 fs. The CHARMM36m force field [23] and TIP3P water model [24] were used. The nonbonding interactions were calculated with a typical cutoff distance of 12 Å, while the long-range electrostatic interactions were enumerated with the Particle Mesh Ewald algorithm [25].

### B. Deep learning analysis

To further understand the molecular structures of different human coronavirus S proteins and the oligomeric state of SARS-CoV-2 S protein, we deployed a custom-built deep learning architecture, a convolutional variational autoencoder (CVAE), to encode the high dimensional protein structures from the MD simulations into lower dimensional latent spaces. The goal of our AI method is to reduce the high dimensionality of the molecular system while preserving the inherent characteristics of the system and learning novel behavior in a latent space that is normally distributed. The direct comparison between the decoded and original input data ensures the accuracy of the latent space representation. This customized CVAE approach has been successfully applied to study the folding pathways of small proteins and structural clustering of biomolecules [26]–[28].

*1) Data preparation:* We used translation and rotation invariant input data for the DL networks. We represented each MD structure by a distance matrix using the *C*_*α*_ atoms of the protein and generated two input datasets for the CVAEs. Input 1 included the matrices of the protomer structures of SARS-CoV-2, SARS-CoV-1, MERS-CoV, and HCoV-HKU1. Input 2 included the matrices of the protomer structure and the chain A of the trimer structure of SARS-CoV-2. For Input 1, as the proteins are different in length, we first performed a multiple sequence alignment of the protomer structures of SARS-CoV-2, SARS-CoV-1, MERS-CoV, and HCoV-HKU1 by Clustal Omega [29]. Based on the alignment, we inserted gaps into the corresponding aligned residue location in the distance matrix and set the distance between gaps to be 0 Å.

The size of each distance matrix after the alignment was 1,342 × 1,342. To reduce the matrix size, we applied a convolution layer with padding of 2 and a 14 × 14 filter with strides of size 7 in both the x and y directions. The size of each resulting matrix became 191 × 191. Finally, after alignment and size reduction we merged a total of 20,000 distance matrices of the four S proteins. An example of a distance matrix following the alignment and size reduction is represented in Fig. S1. For Input 2, as the proteins are of the same length, no alignment was required. The size of each matrix was 959 × 959. We again applied a convolution layer with padding of 1 and a 10 10 filter with strides of size 5 in both the x and y directions to reduce the matrix size. The size of each resulting matrix was also 191 × 191, and we merged a total of 10,000 distance matrices of the protomer and trimer of SARS-CoV-2 S protein.

*2) CVAE implementation:* For each of the two input datasets, we randomly split the aligned matrices into training and validation datasets using the 80/20 ratio and applied a CVAE to capture the important structural features and projected them in the three-dimensional (3-D) latent space for visualization. The encoder network of each CVAE consisted of three convolutional layers and a fully connected layer. We used a 3 × 3 convolution kernel and a stride of 1, 2, and 1 at the three convolutional layers, respectively. We trained each CVAE until the training and validation loss converged. Fig. S2 shows the loss curves along the 150 epochs. The difference between decoded and original images is minimal, suggesting the models were trained successfully (Fig. S3). We then selected representative structures from the clusters in the latent space and visualized them using VMD [30].

All MD simulations and DL analysis were performed on the Summit supercomputer at the Oak Ridge Leadership Computing Facility.

## Acknowledgment

We would like to thank David Bell, Theodore Papamarkou, and Jacob Hinkle for helpful discussions.

This work was performed at the Oak Ridge Leadership Computing Facility (OLCF) of the Oak Ridge National Laboratory (ORNL), which is funded by the Office of Science of the U.S. Department of Energy under Contract No. DE-AC05-00OR22725 and used the Extreme Science and Engineering Discovery Environment (XSEDE) [31] COVID-19 HPC Consortium at the IBM AC922 Summit supercomputer of the OLCF at ORNL through allocation TG-ASC200020.

The research was supported by the U.S. Department of Energy, Office of Science, Office of Advanced Scientific Computing Research, under contract number DE-AC05-00OR22725; the Exascale Computing Project (17-SC-20-SC), a collaborative effort of the U.S. Department of Energy Office of Science and the National Nuclear Security Administration; and in part by the Joint Design of Advanced Computing Solutions for Cancer (JDACS4C) program established by the U.S. Department of Energy (DOE) and the National Cancer Institute (NCI) of the National Institutes of Health. It was performed under the auspices of the U.S. Department of Energy by Argonne National Laboratory under Contract DE-AC02-06-CH11357, Lawrence Livermore National Laboratory under Contract DE-AC52-07NA27344, Los Alamos National Laboratory under Contract DE-AC5206NA25396, Oak Ridge National Laboratory under Contract DE-AC05-00OR22725, and Frederick National Laboratory for Cancer Research under Contract HHSN261200800001E.

## Supplementary Information

**Figure S1.**
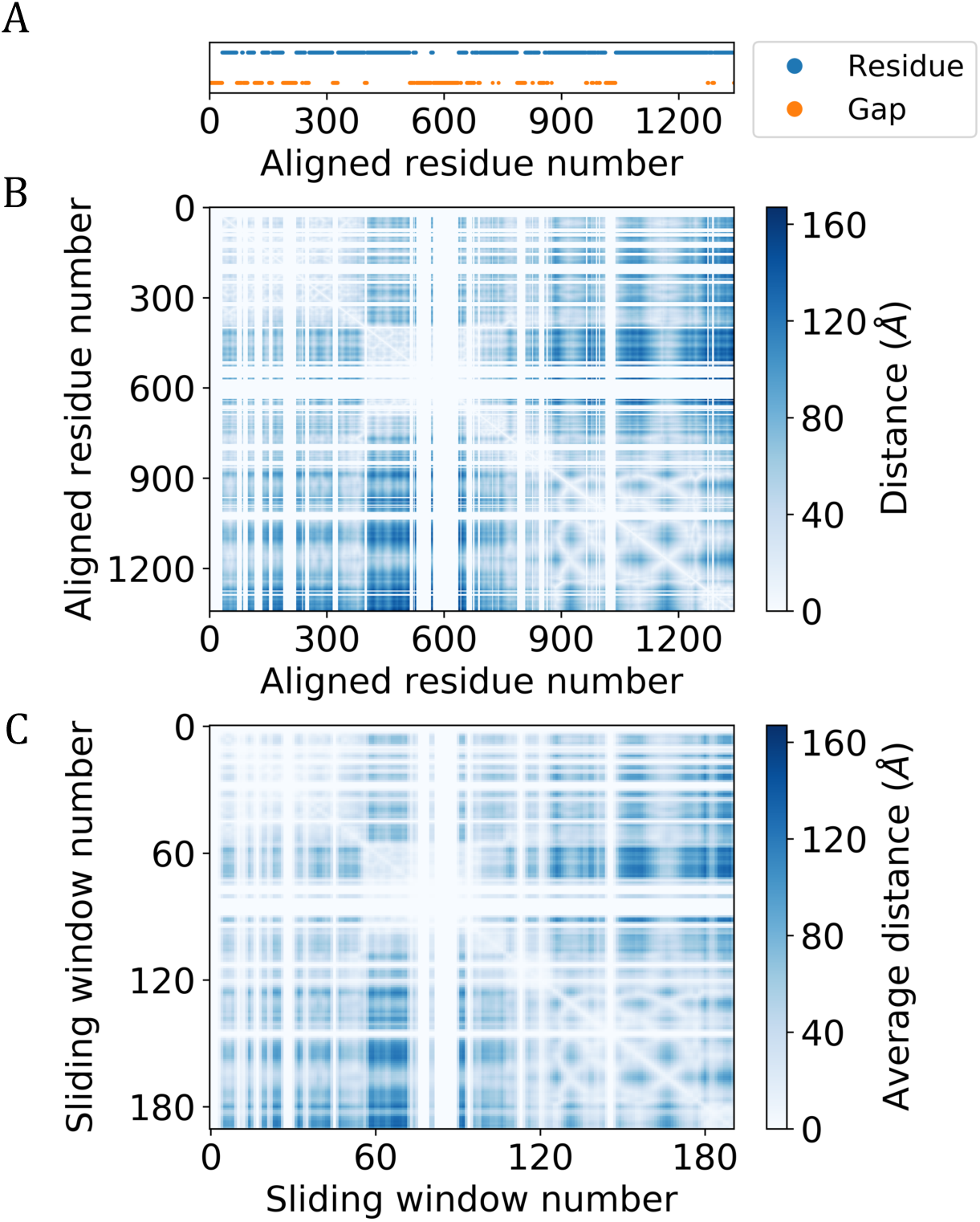
An example distance matrix in Input 1 following the alignment and size reduction. (A) The aligned SARS-CoV-2 S protein sequence with gaps inserted into the corresponding aligned residue location. The 959 residues of PDB 6VSB chain A are colored in blue, and the gaps are in orange. (B) The distance matrix with gaps inserted according to the aligned sequence shown in (A). The distance between gaps was set to be 0 Å. (C) The distance matrix shown in (B) after the size reduction by applying a convolution layer. Note that major patterns are preserved in the matrix of reduced dimensions. (More details are in Methods.)

**Figure S2.**
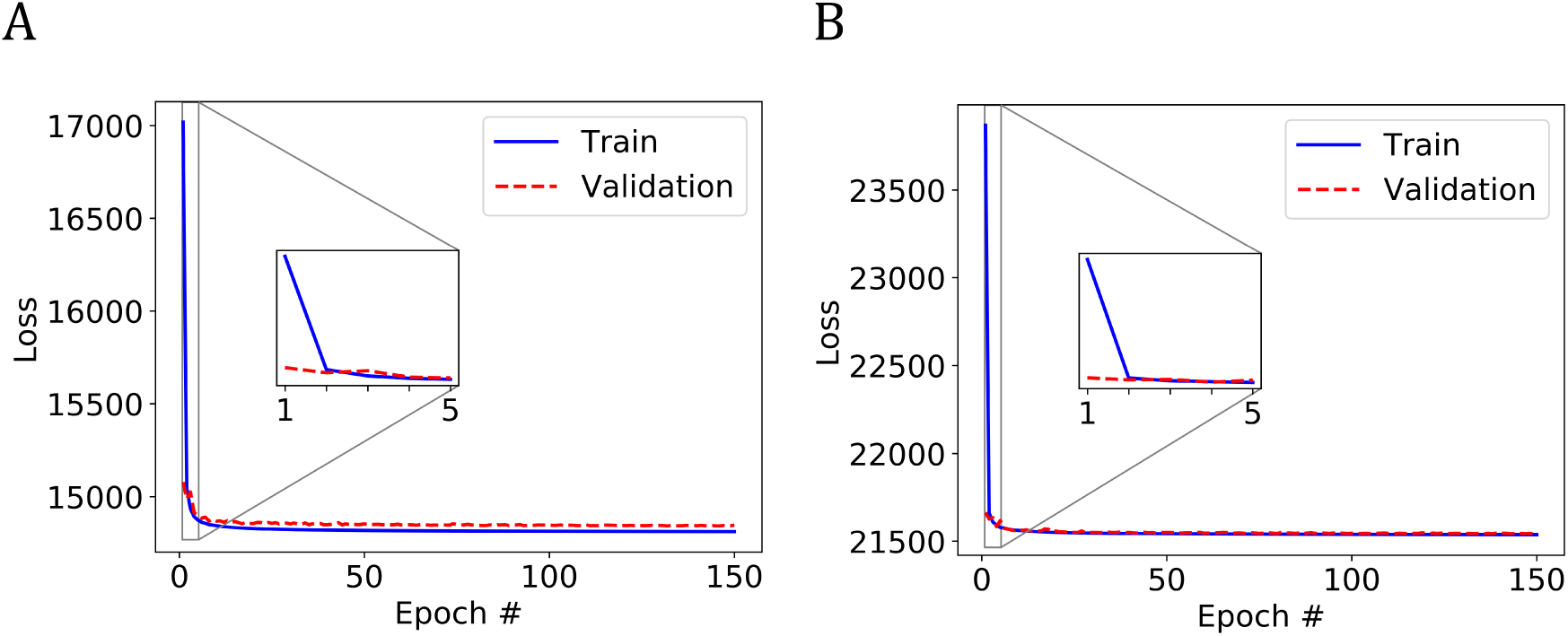
Training and validation loss curves. The loss curves when training the CVAEs on (A) Input 1 and (B) Input 2 along the 150 epochs. The insets show the loss curves in the first 5 epochs.

**Figure S3.**
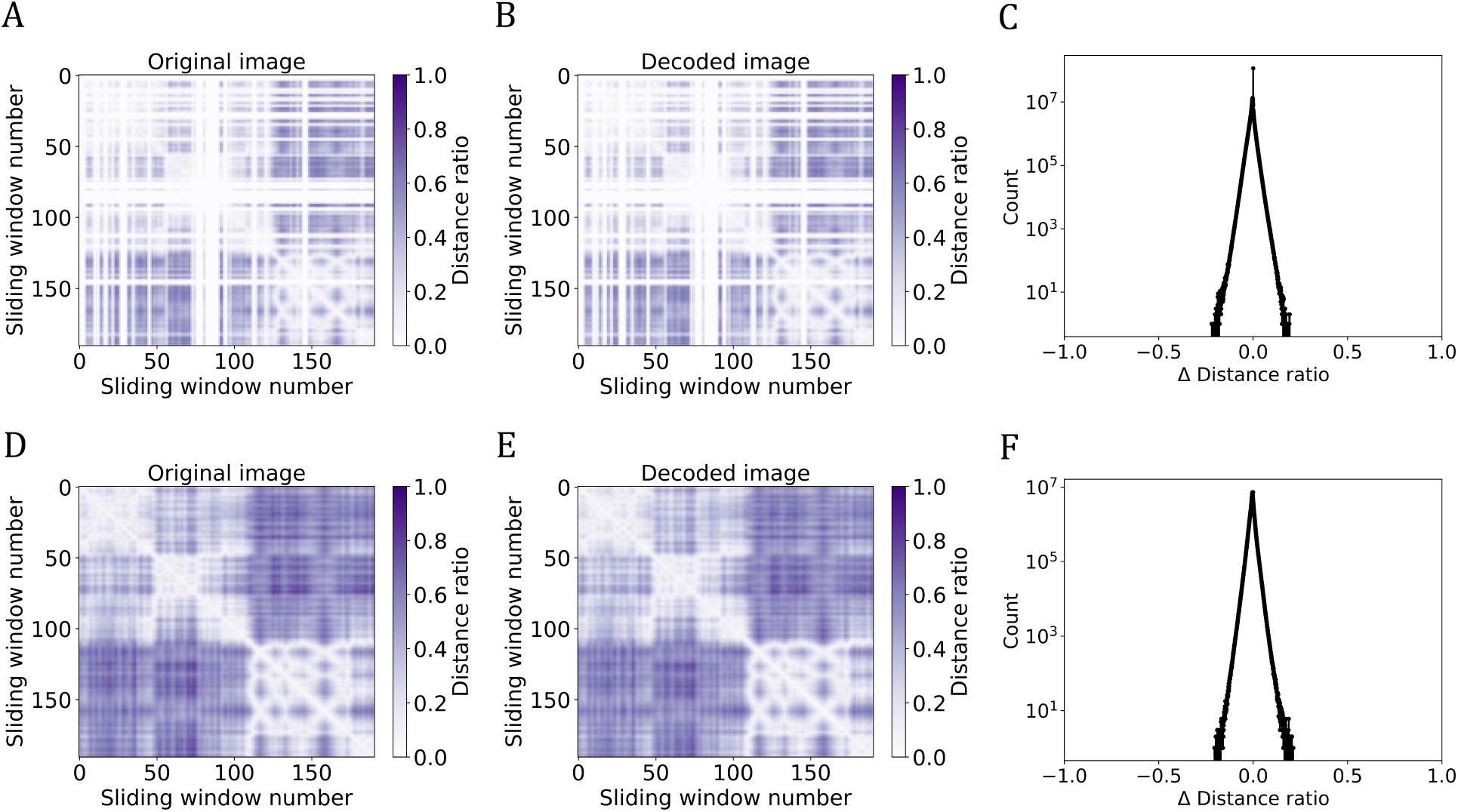
CVAE model assessment. Assessment of the model trained on (A-C) Input 1 and (D-F) Input 2 after 150 epochs. (A, D) An example original image. (B, E) The decoded image using the trained models. (C, F) Enumerated difference between the decoded and original images in the corresponding input dataset. Note that the y-axis is in log scale.

**Figure S4.**
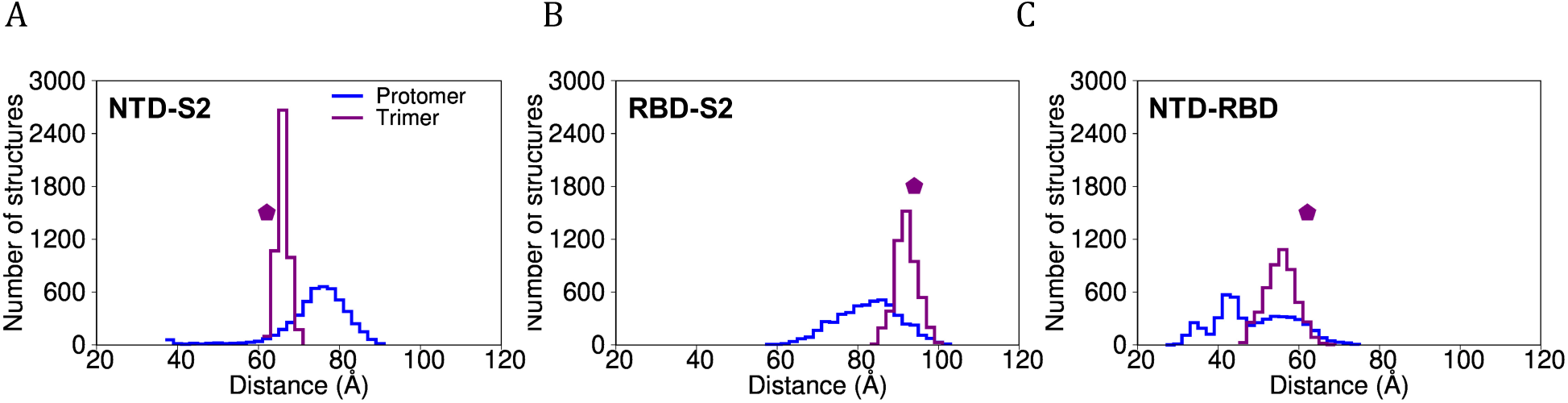
Distribution of the interdomain distances of the protomer and chain A of the trimer of SARS-CoV-2 S protein. The distribution of the center-of-mass distance between the backbone atoms of the residues that constitute (A) NTD and S2 domains, (B) RBD and S2 domains, and (C) NTD and RBD domains. The distribution of the protomer is colored in blue and that of the trimer in purple. The distances calculated from the cryo-EM structure (PDB 6VSB) are marked by a pentagon.

